# Computational Tools for the Identification and Interpretation of Sequence Motifs in Immunopeptidomes

**DOI:** 10.1101/210336

**Authors:** Bruno Alvarez, Carolina Barra, Morten Nielsen, Massimo Andreatta

## Abstract

Recent advances in proteomics and mass-spectrometry have widely expanded the detectable peptide repertoire presented by major histocompatibility complex (MHC) molecules on the cell surface, collectively known as the immunopeptidome. Finely characterizing the immunopeptidome brings about important basic insights into the mechanisms of antigen presentation, but can also reveal promising targets for vaccine development and cancer immunotherapy. In this report, we describe a number of practical and efficient approaches to analyze immunopeptidomics data, discussing the identification of meaningful sequence motifs in various scenarios and considering current limitations. We address the issue of filtering false hits and contaminants, and the problem of motif deconvolution in cell lines expressing multiple MHC alleles, both for the MHC class I and class II systems. Finally, we demonstrate how machine learning can be readily employed by non-expert users to generate accurate prediction models directly from mass-spectrometry eluted ligand data sets.

## Introduction

The comprehensive set of peptides presented on the cell surface by MHC molecules, collectively referred to as the immunopeptidome, represents a unique fingerprint of the health of a cell. T lymphocytes routinely scan this pool of MHC-associated peptides, and can help eliminating infected or cancerous cells that present abnormal peptides on their surface. MHC class I molecules mainly bind peptides derived from intracellular pathogens (such as viruses and some bacteria) and present them to cytotoxic T lymphocytes; MHC class II epitopes are mainly derived from extracellular proteins and are presented to T-helper lymphocytes.

Recent technological advances in the field of mass spectrometry (MS) have brought about a revolution in the study of immunopeptidomes (reviewed in ref. [1]), with several thousands of peptides that can be detected in a single experiment. Large data sets of naturally presented peptides have been beneficial to define more accurately the rules of peptide-MHC binding [2–4] but have also a tremendous potential in defining pathogen-derived T cell epitopes [5,6] and neo-epitopes unique to cancerous cells [7,8]. Part of the appeal of MS-based approaches is that they do not require prior knowledge of MHC motifs, and there is no human intervention in defining a library of candidate sequences to be tested. Therefore, MS provides a large but relatively unbiased sampling of the population of processed and presented peptides available for T cell recognition [3].

In most MS-based pipelines, spectra from eluted peptides are matched against a reference database of natural proteins using algorithms like MaxQuant [9] or PEAKS [10,11], and filtered against a decoy database to limit the false discovery rate (FDR). Strict FDR filters (typically in the order of 1%) should ensure that most spectra are correctly assigned to *bona fide* ligands, but often leads to discarding a large portion of the spectra. Several approaches have been proposed to increase the yield of spectral assignment. For example, Mascot Percolator performs machine learning on high-confidence matches to rescore database search results for lower-confidence peptides [12]. Instead of matching spectra to an entire protein database, SpectMHC constructs reduced, targeted databases of potential MHC ligands, effectively reducing the amount of spurious decoy hits [13]. A sizable portion of the unassigned spectra may also be explained by proteasome-generated spliced peptides, which would require the inclusion of spliced variants in the target database [14].

After spectral assignment to amino acid sequences, peptides must often be aligned and/or clustered to extract meaningful sequence motifs of antigen presentation. The analysis protocols here will generally differ depending on the type of receptor (MHC I vs. MHC class II) and type of sample used (cellular versus soluble MHC molecules and mono-vs. poly-allelic cell lines). MHC I ligands have a limited range of lengths, typically 8 to 11 amino acids long, and are characterized by very conserved amino acid preferences at the positions interacting with the MHC binding groove (anchor positions). On the other hand, MHC II ligands are normally longer, with only a portion, the binding core, directly interacting with the MHC groove [15]; in this case a more sophisticated alignment process is needed to extract conserved binding preferences. In transgenic cells expressing a single MHC molecule (mono-allelic), only one specificity is expected to be present in the data and motif identification is relatively straightforward. Conversely, unmodified cells will naturally present peptides bound to multiple MHC alleles (up to six for HLA class I), with generally different binding preferences; in this case, the multiple specificities contained in the data must be deconvoluted, either by assigning MHC restriction with predictive methods, or by unsupervised clustering.

A popular tool for the unsupervised identification of sequence motifs in immunopeptidomes is GibbsCluster [16,17], a web-based and downloadable method that has been included into numerous pipelines for the deconvolution of ligand motifs in the MHC class I [18–21] and MHC class II [22–24] systems. The GibbsCluster algorithm takes as input a list of peptide sequences (potentially of variable length), and uses a heuristic search to group them into information-rich groups. Besides the sequence motif defining each group, additional properties such as the ligand length distribution of each cluster can be analyzed. A similar method, MixMHCp [2,25], has shown performance comparable to GibbsCluster, with the limitation that it can only handle peptides of uniform length. A useful feature of GibbsCluster is the “trash cluster”, a check on internal motif consistency that can filter out outliers that cannot be assigned to any clusters. In the context of MS eluted ligand data, spurious data points can originate both from LC-MS/MS contaminants and from erroneous spectral matches. As a noise filter, GibbsCluster can be beneficial also for mono-allelic data sets where no motif deconvolution is required.

While sequence motifs are generated by GibbsCluster in an unsupervised manner, the method cannot directly assign the MHC restriction of each ligand; this must be done by comparing the unsupervised motifs with published binding motifs of the MHC molecules in the sample [26]. While this comparative approach is in most cases feasible for human MHC, whose most prevalent alleles have been well characterized and documented, it will fall short for samples containing uncharacterized specificities. Aiming to overcome this limitation, Bassani-Sternberg et al. [25] suggested a strategy for automatic, unbiased annotation of MHC restriction by comparing motifs detected in multiple data sets with known haplotypes. Exploiting the co-occurrence of MHC alleles across different data sets, they were able to assign motifs to individual alleles without relying on *a priori* assumptions on their binding specificity, also for alleles without previously documented ligands.

Over the past decades, many efforts have been dedicated to the development of computational methods for the prediction of peptide binding to MHC class I molecules. Most of these T-cell epitope prediction methods have been traditionally trained solely on *in-vitro* data of peptide-MHC binding affinity. Although peptide-MHC affinity is arguably the most selective step in antigen presentation, other factors influence the likelihood of a peptide being presented on the cell surface for T-cell recognition [27,28]. *In-vitro* binding affinity data does not address the fact that antigen presentation is a complex, integrative physiological process that combines antigen processing, transport and binding affinity/stability of the peptide-MHC complex. Finally, *in-vitro* data fails to reflect any peptide length preference of different MHC-I alleles. Because naturally eluted ligands incorporate information about these additional properties of antigen presentation, large MS-derived sets of peptides can potentially enable the generation of more accurate prediction models. Recent studies have suggested that models trained on MHC class I ligand data outperform binding affinity-based predictors when it comes to identification of eluted ligands and T cell epitopes, both in an allele-specific setting [2,3] as well as with pan-allelic coverage [4]. Generic tools for machine learning from peptide sequences such as NNAlign [29,30] can be applied to individual MS data sets to generate custom-made prediction models, which can in turn be employed for further downstream analyses of the immunopeptidome.

The rapidly expanding collection of naturally eluted ligands revealed by MS and the analysis toolkits developed in its wake hold great promise in understanding the structure of the immunopeptidome and the rules of antigen presentation. However, because of the complexities inherent to MS eluted ligand data, it is not a trivial task to analyze and interpret the information these data sets contain. In this report, we seek to address some common issues and describe strategies to analyze MS ligand data and derive sequence motifs in the various scenarios outlined above (MHCI vs. MHCII; mono-allelic vs. poly-allelic cell lines), with guidelines and examples on publicly available datasets.

## MHC class I; mono-allelic cells

In a recent publication, Abelin et al. [3] described the development of transgenic cells that express a single human MHC class I allele (HLA), and used them to generate a large set of MHC ligands covering 16 HLA class I alleles. There are obvious advantages in using mono-allelic cells to characterize MHC ligands: firstly, no deconvolution/clustering is required to define motifs at the single-allele resolution; secondly, the assignment of individual peptides to their allele does not have to depend on binding predictions or prior knowledge of the motifs. Apart from technical difficulties in the cell generation, a possible drawback is that the relative level of expression of different MHC alleles in a given cell, and the amount of ligands they present, is lost in a mono-allelic setting. The amount of ligands presented by different alleles may also depend on competition between MHC molecules, where the newly available digested peptides from an unfolding antigen fragment would presumably be captured by MHCs with the highest affinity [31].

Although most software for MS spectra mapping uses a strict false discovery rate (FDR) threshold, incorrect ligands may still be present among the matches that pass the FDR check. These may consist of common contaminants such as keratin or histone proteins, as well as residual peptides from previous runs of the LC-MS/MS instruments used for sample preparation [32,33]. GibbsCluster is a useful tool to detect and remove such contaminants and false hits. For each allele in the Abelin data set [3], we applied GibbsCluster-2.0 with default preset options for “MHC class I ligands of length 8-13”, specifying a single cluster. Between 0.4% and 16% of the peptides (mean 4%) of length 8 to 13 were inconsistent with the motif identified by GibbsCluster-2.0 and were removed by the program as noise. While distinct motifs can be discerned before trash cluster filtering (see three representative alleles in Figure 1A), the post-filtering motifs have higher information content and more well-defined anchor residues (Figure 1B). Peptides in the “trash cluster” may sometimes hint at the origin of the contamination: for example, the observation of terminal Arginine/Lysine preferences at the C-terminus in several of the 16 alleles points towards tryptic peptides polluting the mixtures (Supplementary Figure S1). The ligands in the Abelin data set have in general very good correspondence to known MHC binding preferences, with an average NetMHCpan-3.0 percentile rank [34] well below 1% for most alleles (Figure 1C, red boxplots). In contrast, peptides in the trash cluster match very poorly the preferences of their MHC and are assigned high NetMHCpan rank scores (Figure 1C, blue boxplots).

**Figure 1.**
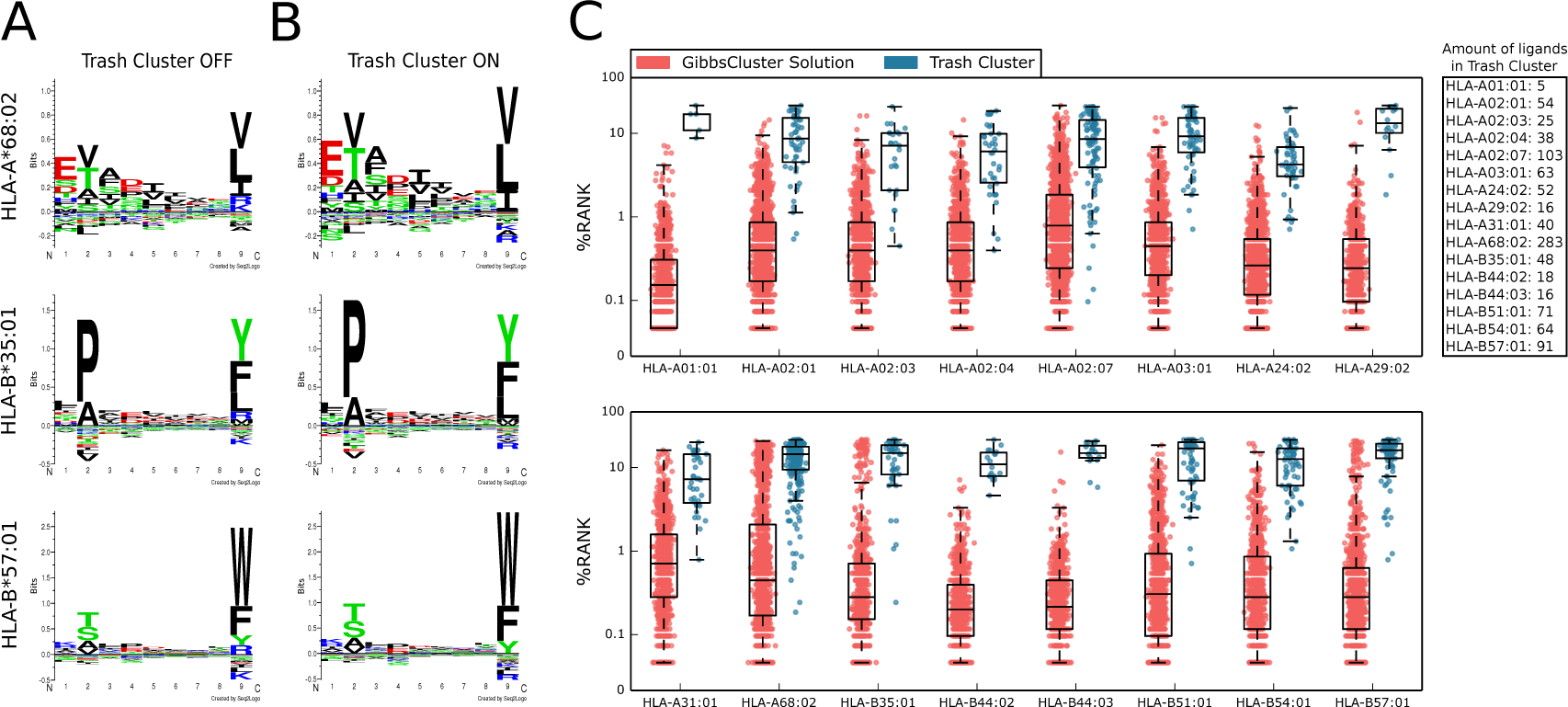
Visualizing motifs and removing contaminants with GibbsCluster. A) Sequence motifs of three representative alleles before trash cluster filtering and **B)** after filtering. The post-filtering motifs have higher information content and lack the putative K/R contamination at P9. **C)** Distribution of NetMHCpan-3.0 percentile rank scores for peptides in the main cluster (red) and in the trash cluster (blue).

## MHC class I; poly-allelic cells

Unmodified antigen-presenting cells will generally express up to six different MHC class I alleles (two each for HLA-A, HLA-B and HLA-C). The immunopeptidome of these cells therefore consists of multiple specificities mixed together, where the global haplotype is known but the restriction of each individual ligand is unknown. For example, Bassani-Sternberg et al. [18] described the LC-MS/MS analysis of peptides eluted from seven different cancer cell lines and primary cells, which had been HLA-typed at high resolution, and demonstrated how the GibbsCluster approach could be used to deconvolute the individual peptide restrictions. Here we illustrate the application of GibbsCluster to one of the cell lines from the Bassani-Sternberg study, HCC1143, which expresses the five alleles HLA-A*31:01, HLA-B*35:08, HLA-B*37:01, HLA-C*04:01, HLA-C*06:02.

GibbsCluster finds an optimal solution of four clusters, with a close correspondence to all but one of the HC1143 alleles (Figure 2), failing to separate HLA-C*04:01 ligands. HLA-C molecules have low expression levels and rather degenerate binding preferences [25,35], making the deconvolution of their motifs more challenging. The motifs determined by unsupervised clustering show a remarkable correspondence with the binding preferences predicted by NetMHCpan [34], with, in some cases, additional secondary anchors (e.g. a positively charged P5 for HLA-B*37:01) that may contribute to enhance peptide-MHC complex stability. The sizes of the clusters give an indication of the relative level of expression of the different alleles, with the largest group corresponding to the homozygous HLA-A*31:01 (1253 peptides), followed by the two HLA-B alleles (610 and 460 peptides respectively) and by the lowly-expressed HLA-C*06:02 (409 peptides). Finally, 45 peptides were collected by the trash cluster. Interestingly, for six out of seven cell lines in the Bassani-Sternberg data set, we noted a C-terminal enrichment for Arginine/Lysine in peptide discarded in the trash cluster (Supplementary Figure S2). A similar observation was made for the Abelin data set discussed previously, and hints that residual peptides derived from trypsin digestion may often be present in the LC column.

**Figure 2.**
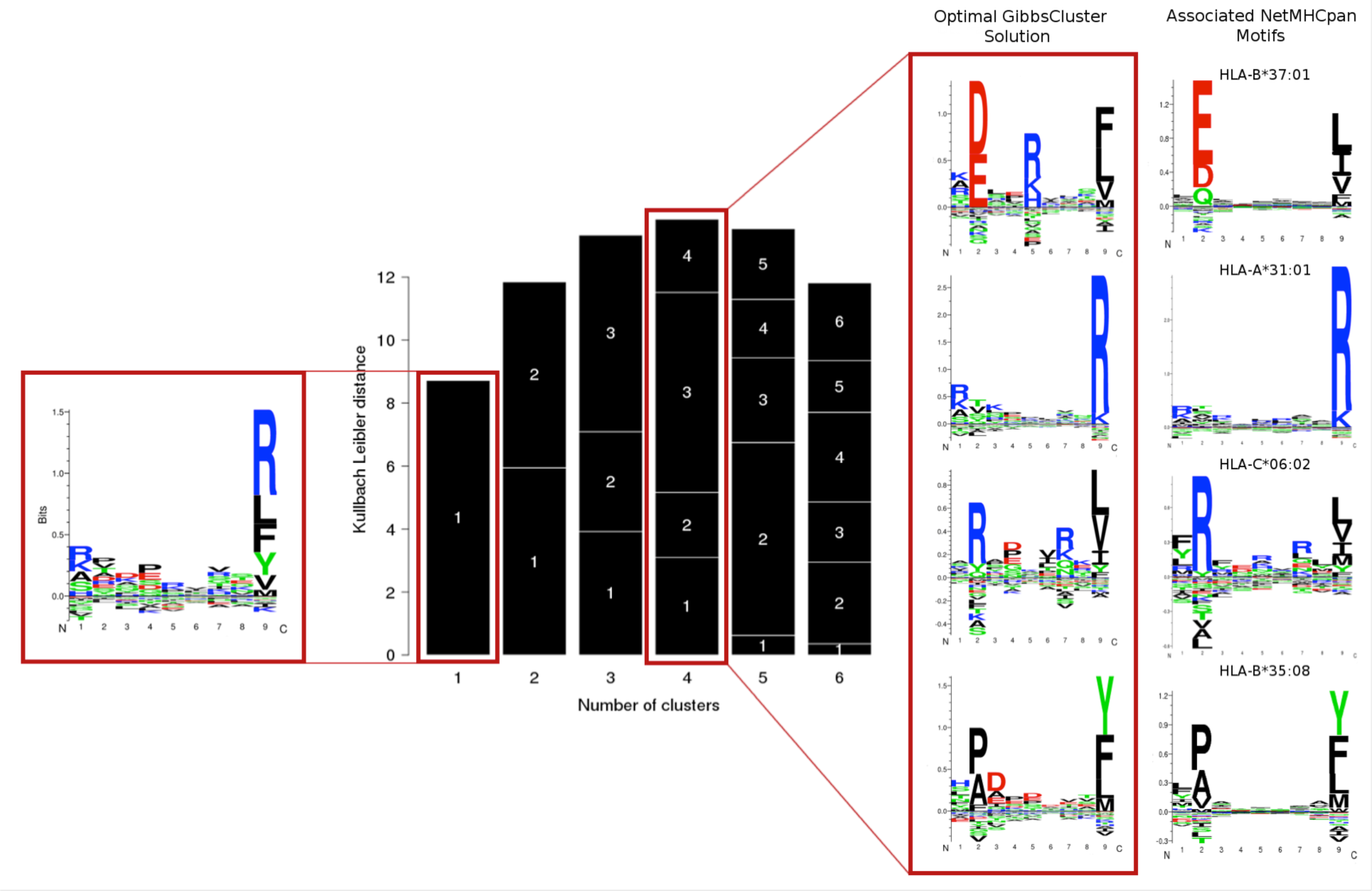
Clustering results for the HCC1143 cell line. The single cluster solution **(left)** is a mixture of multiple specificities, dominated by the most abundant alleles. The solution with highest information content corresponds to four clusters, with motifs highlighted in the red box **(center)**. The motifs identified by unsupervised clustering show a remarkable correspondence with those predicted by NetMHCpan-3.0 **(right)**. The GibbsCluster method was run using the default preset parameters for “MHC class I ligands of length 8-13”, except for the number of iterations which was set to 100 (slower but more accurate), and number of groups, which was allowed to vary between 1 and 6.

## MHC class II, mono-allelic cells

Analyzing MHC class II binding data is for many reasons more complex compared to MHC class I. First and foremost, the HLA class II binding groove is open at both ends, accommodating peptides of a wide range of length by letting them protrude at either terminus of the nonamer binding core. Sophisticated alignment methods are therefore required to identify the conserved binding preferences of MHC class II molecules [36,37]. Secondly, the binding motifs for MHC class II are in general more degenerate compared to the highly conserved MHC class I motifs [38,39]. These observations make the analysis and interpretation of MHC class II binding data, including MS ligands, highly challenging.

In a recent paper by Ooi et al. [40], MS eluted ligand data were used to investigate how patients expressing different HLA class II alleles have different susceptibility to autoimmune diseases. To characterize the specificity for each allele, they generated transgenic mice bearing the human HLA-DR1 MHC class II allele. On these data, we illustrate how the GibbsCluster method can be used to identify the binding motif of MHC class II molecules from mono-allelic MS ligand data and at the same time remove potential outliers. The 5740 non-redundant raw eluted peptide sequences were uploaded to the GibbsCluster web server, setting the recommended preset parameters for MHC class II peptides, except for the number of iterations per sequence per temperature step (set to 100) and the number of temperature steps (set to 50); these parameters entail a slower, but more accurate, motif search. The method recovered the binding motif for allele HLA-DRB1*01:01, with strong amino acid preferences at anchor residues at P1, P4, P6 and P9 (Figure 3A). These preferences were observed both without (Left logo, figure 3A) or with a trash cluster activated (Right logo, figure 3A). By activating the trash cluster option with a threshold of 2, 179 peptides (3% of data) were removed, and the logo showed a 20% increase in information content (Right logo, figure 3A).

**Figure 3.**
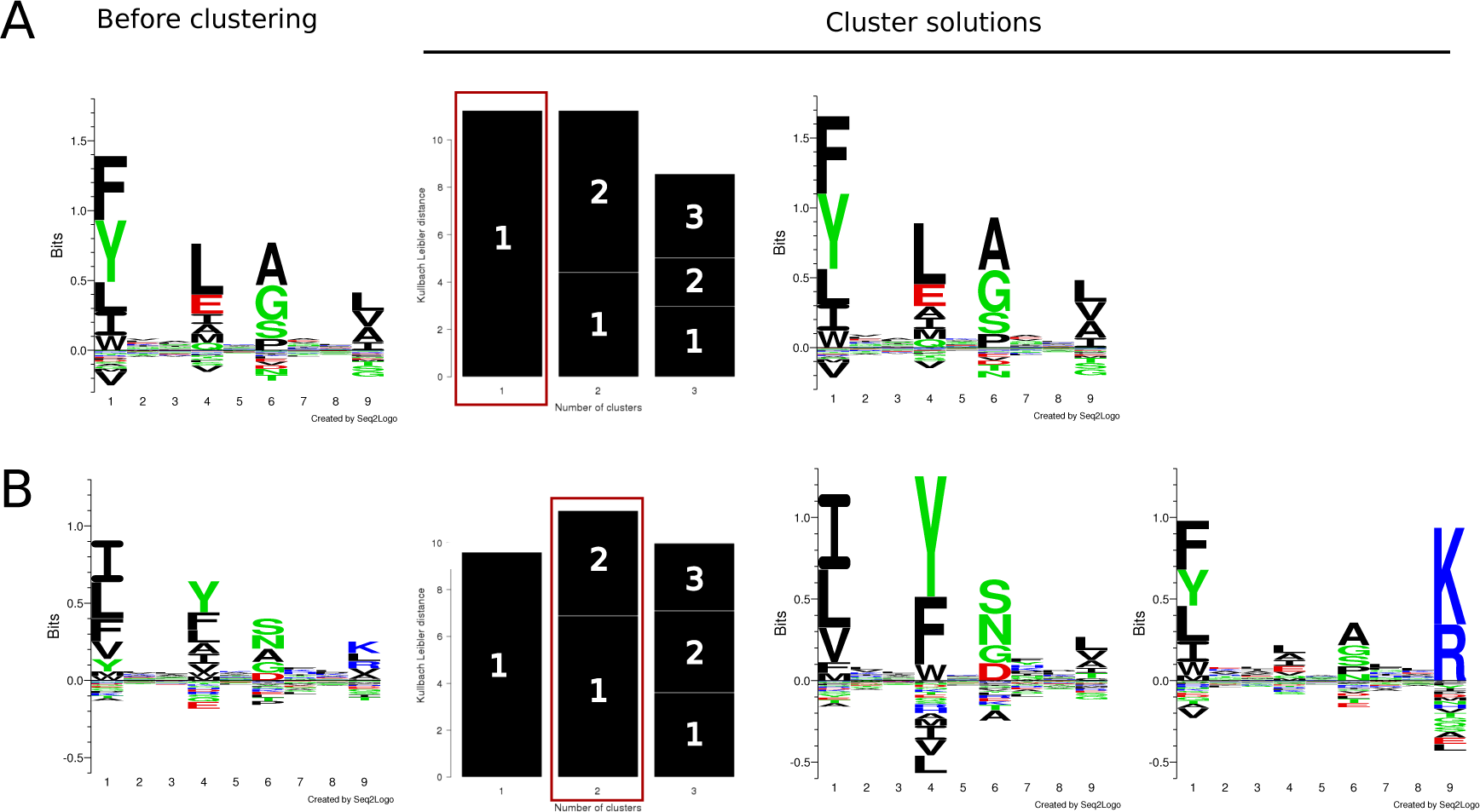
Sequence motifs identified by GibbsCluster-2.0 on MHC class II ligand data. The method identifies distinct amino acid preferences at the anchor positions P1, P4, P6 and P9 both without (left panels) and with (right panels) the trash cluster activated. **(A)** Visualizing the motif and removing outliers from the mono-allelic human-DR1 mouse-transfected cell lines. **(B)** Motif identification on mixed allelic data of DR15-DR51-EBV transformed cell lines.

## MHC class II, poly-allelic cells

Another data set obtained from the Ooi et al. study [40] consists of peptides eluted from HW09013 cells that express the HLA-DR15/DR51 class II alleles. On this poly-allelic data set of MS eluted ligands, we set out to demonstrate how the GibbsCluster can be used to separate multiple specificities in MHC class II ligand data. The set of 2782 unique eluted peptides was submitted to GibbsCluster, using the recommended preset parameters for MHC class II and allowing the program to search up to three clusters. The unfiltered, single-cluster solution shows a motif with the correct P1, P4, P6 and P9 anchor positions, but with low information content and preferences that are a mixture of the two alleles in the sample (Figure 3B, left logo). Activating the trash cluster with a threshold of 2, the maximum information content is observed for the solution with two clusters (Figure 3B, bar plot). The amino acid preferences identified by GibbsCluster resemble previously published motifs derived from binding affinity data for HLA-DRB1*15:01 and HLA-DRB5*01:01 [29,41], and closely overlap with the global peptidome of DR15/51 characterized in a recent study [42]. Specifically, cluster 1 was composed of 1610 peptides (57.9%) and its motif resembles the HLA-DR15 binding preferences; cluster 2 comprised 1050 peptides (37.7%) and corresponds to the HLA-DR51 alleles; 122 peptides (4.4%) did not match to either group and were collected by the trash cluster.

In order to validate the solutions generated by GibbsCluster, we examined the composition of the clusters in terms of binding potential predicted by NetMHCIIpan-3.1 [43]. Both for the mono-allelic DR1 and poly-allelic DR15/51 serotypes discussed above, we obtained predicted percentile rank scores for all peptides in the cluster solutions and in their relative trash cluster (Figure 4). The predicted median rank score for HLA-DRB1*01:01 in the DR1 cluster was 4% (first quartile (Q1)=0.9, third quartile (Q3)=12), whereas the trash cluster had a median rank score of 41% (Q1=20.5, Q3=75). In the poly-allelic data, cluster 1 was associated with HLA-DRB1*15:01, and showed a median rank score of 13% (Q1=5, Q3=30); cluster 2 was associated to HLA-DRB5*01:01 and obtained a median rank score of 4% (Q1=1.1, Q3=11); peptides in the trash cluster were evaluated against both alleles, assigning the best rank of the two, which resulted in an average rank score of 41% (Q1=23, Q3=75) (Figure 4). Overall, the NetMHCIIpan percentile score distributions suggest that the trash cluster could successfully collect peptides with very poor correspondence to the known preferences of the MHC class II molecules, and that probably derived either from incorrect spectral matches or from contaminants. The relatively high predicted rank values for the peptides mapped to the HLA-DRB1*15:01 cluster further suggest that the binding motif for this molecule predicted by NetMHCIIpan-3.1, which was trained on binding affinity data, shared a rather weak overlap with the binding motif contained within the MS ligand data. This observation underlines the high potential of MS ligand data to complement our knowledge on peptide characteristics required for MHC antigen presentation, as previously remarked for MHC class I [3,4].

**Figure 4.**
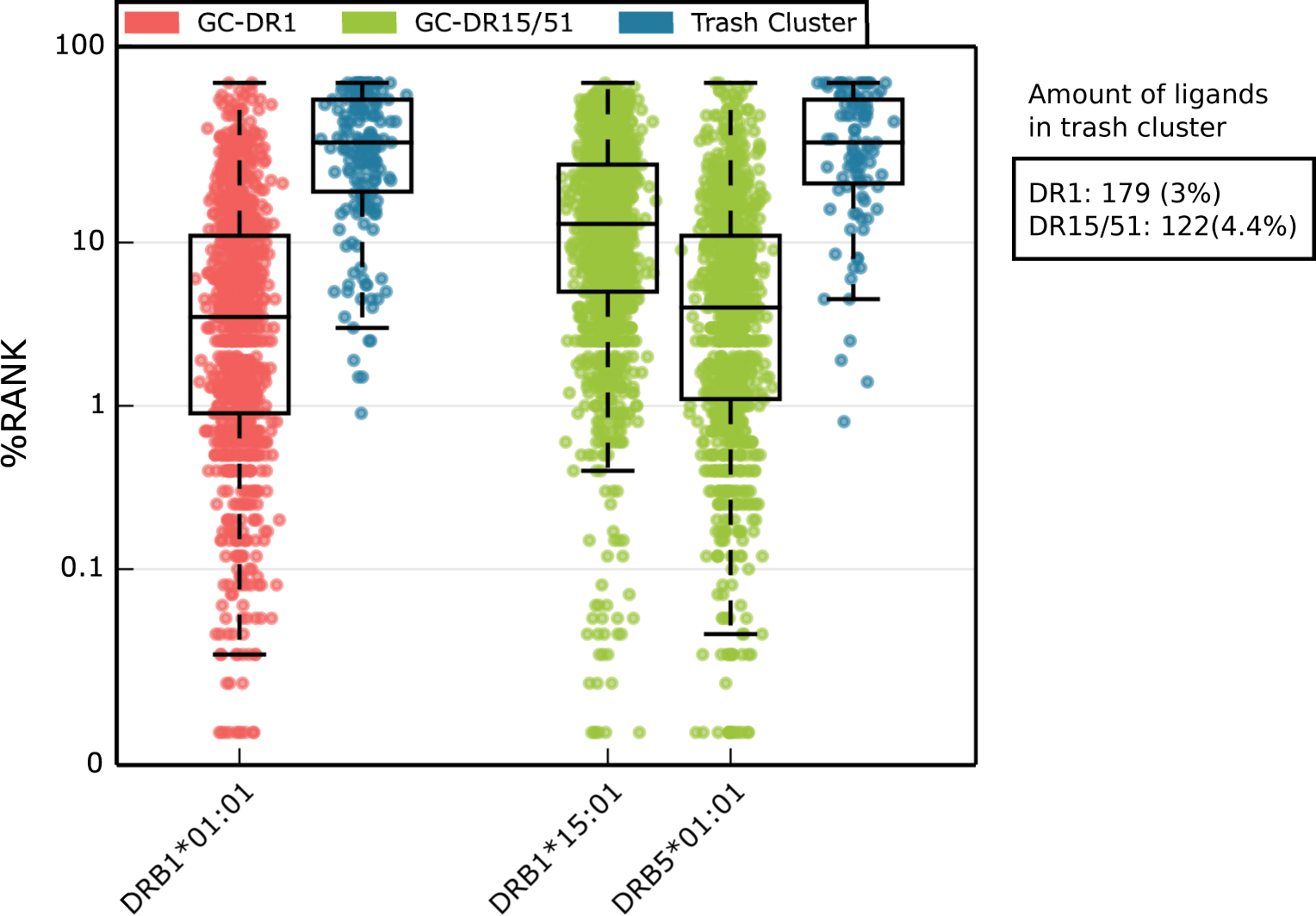
NetMHCIIpan percentile rank score for GibbsCluster solutions in the DR1 and DR15/51 data sets. Percentile rank scores were predicted by netMHCIIpan-3.1 for each GibbsCluster group with matching alleles present in MS data samples. In the case of the mixed allele dataset DR15/51, peptides in the trash cluster were scored by NetMHCIIpan to both DRB1*15:01 and DRB5*01:01, selecting the lowest rank score of the two.

## Generating prediction models from MS ligand data

The approaches described so far in this report are mainly concerned with extracting and visualizing meaningful patterns within complex, often noisy, mixtures of peptides sequences. A further step is the generalization of the motifs identified in the data at hand, by constructing prediction models. Machine learning algorithms such as NNAlign [30], when provided with training examples suitably labeled (e.g. ligands vs. non-ligands), can be instructed to automatically learn the features that distinguish positive from negative examples. Such models can then be applied on external data sets to discover more occurrences of the patterns learned on the training data. In the context of peptide-MHC interactions, a good prediction model should have the ability to capture the binding preferences contained in the training data, both in terms of sequence motifs and peptide length distribution. In the next two sections, we illustrate some simple examples of prediction models directly constructed from MHC class I and class II eluted ligands.

### MHC Class I prediction model

As an example application, we continue with the Abelin ligand elution dataset previously analyzed and filtered using GibbsCluster-2.0 (Figure 1). For each of the representative alleles HLA-A*68:02, HLA-B*35:01 and HLA-B*57:01, we prepared a training set consisting of post-filtering ligands (positive instances) and random natural peptides (negative instances). Positive instances were labeled with a target value of 1, negatives with a target value of 0. In line with earlier work [4], the amount of random negatives was imposed to be the same for each length 8 to 13, and corresponded for each length to five times the amount of positives for the most abundant peptide length. This uniform length distribution of the random negatives was adopted as a background against with machine learning can be employed to learn the amino acid and length preference of the natural binders.

On each of the three data sets, we trained a prediction model with the NNAlign-2.0 webserver, using the recommended preset options for MHC class I ligands of variable length. In a cross-validation experiment, the three models returned an area under the ROC curve (AUC) of 0.961, 0.984 and 0.979, respectively. In order to derive the amino acid and peptide length preferences learned by the model, we used it to evaluate a large set of 900,000 random natural peptides with a flat length distribution, and extracted the top 1% scoring peptides. The composition of these high-scoring peptides should reflect the main preferences identified by the method to distinguish positive from negative instances. Indeed, the binding motif drawn from the top 1% peptides closely reflects the amino acid preferences of the training data (Figure 5A-B). Moreover, all three methods could capture the preference for 9mer peptides over other peptide lengths; 10mers were moderately allowed, 8mers and 11mers were observed more infrequently (Figure 5C).

**Figure 5.**
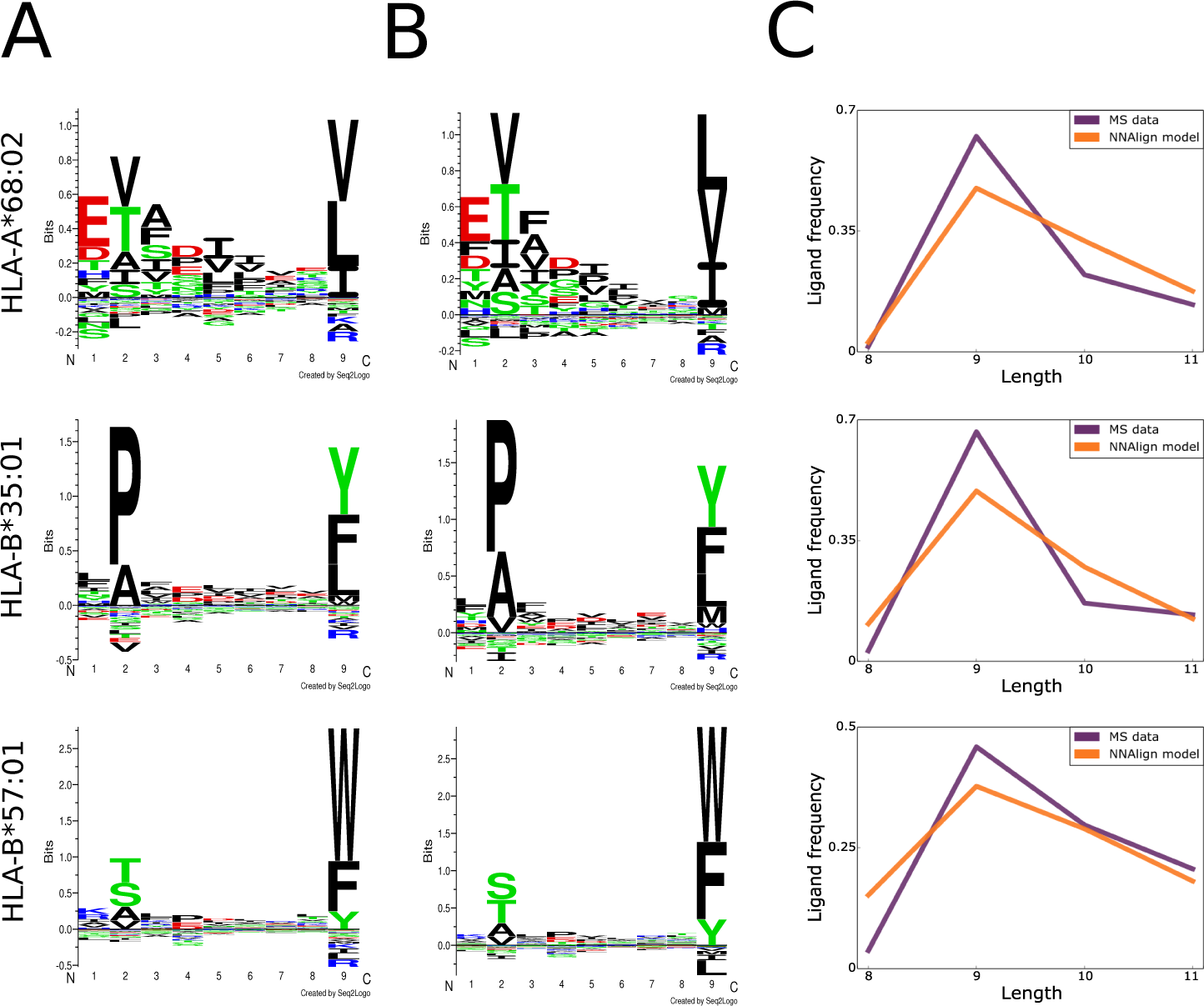
Generating prediction models from MS ligand data. **A)**Sequence motifs of the training data for three MHC class I alleles, aligned and filtered by GibbsCluster; **B)** Sequence motifs captured by NNAlign-2.0; **C)** Ligand length preferences in the training MS data compared to length preferences learned by the NNAlign model.

### MHC Class II prediction model

To illustrate how the NNAlign framework can be used to construct MHC class II prediction models, we go back to the DR1 and DR15/51 data sets from Ooi et al. [40] previously filtered and clustered with GibbsCluster (Figure 3). To enrich the positive instances with artificial negative examples, a set of natural random negatives of length 11 to 19 amino acids was added to each eluted ligands data set. Positive instances were labeled with a target value of 1, negatives with a target value of 0. Similarly to the training set preparation described above for MHC class I, the amount of random negatives for each length corresponded to five times the amount of positives for the most abundant peptide length. For each of the three specificities deconvoluted by GibbsCluster in the DR1 and DR15/51 cells, we applied NNAlign-2.0 to generate a prediction model, using the preset parameters for MHC class II recommended by the NNAlign server. For the mono-allelic DR1 serotype, all ligands except those removed by the trash cluster were used to train a model. For the DR15/51 cells, for which the clustering analysis revealed two separate specificities, we generated a separate model from the ligands contained in each of the two clusters.

The models revealed high internal consistency, with cross-validated performance of AUC=0.952, 0.974 and 0.952 for respectively. NNAlign automatically generates a matrix (and logo) representation of the motif learned by the method, constructed from the top 1% scoring predictions from a large set of random natural peptides. We may compare the motifs learned by NNAlign to: *i)* the binding preferences in the MS training data, identified by GibbsCluster; *ii)* the GibbsCluster motifs identified in tetramer-validated epitopes extracted from the IEDB for the three DR molecules; *iii)* the binding preferences predicted by NetMHCIIpan-3.1 for these DR molecules. In general, the motifs learned by the NNAlign models share a remarkable overall correspondence to the preferences found by GibbsCluster for the MS ligand data, with similar amino acid enrichments at the anchor positions P1, P4 and P6, as well as the strong P9 for the DR51-associated ligands (Figure 6, first and second columns). Likewise, the binding motifs constructed from the rather small amount of tetramer-validated epitopes obtained from the Immune Epitope Database (IEDB) [44] for the three DR molecules (231 for HLA-DRB1*01:01, 129 for HLA-DRB1*15:01, 73 for HLA-DRB5*01:01) correspond well with the motifs of the NNAlign models, and the MS ligand data (Figure 6, third column). In contrast, the logos derived from *in-vitro* binding affinity data (NetMHCIIpan) in all cases show substantial differences to both the MS- and epitope-derived motifs (Figure 6, fourth column). These discrepancies are most pronounced for HLA-DRB1*15:01, where the NetMHCIIpan motif has weakly defined preferences at the anchor residues, and an enrichment of arginine (R) throughout the binding motif; a preference that is completely absent from the MS and epitope-derived motif. Another, more subtle difference is the enrichment of glutamic acid (E) at P4 in the MS and epitope motifs for HLA-DRB1*01:01; this preference is absent in the NetMHCIIpan motif. Finally, NetMHCIIpan displays a preference for R/K at position P8 for HLA-DRB5*01:01; this anchor is completely absent in the motif derived from MS and tetramer-validated epitope data. Taken together, these results show that ligand elution is a stronger correlate of epitope presentation than peptide-MHC binding affinity, suggesting that epitope prediction models may greatly benefit from incorporating MS eluted ligand data.

**Figure 6.**
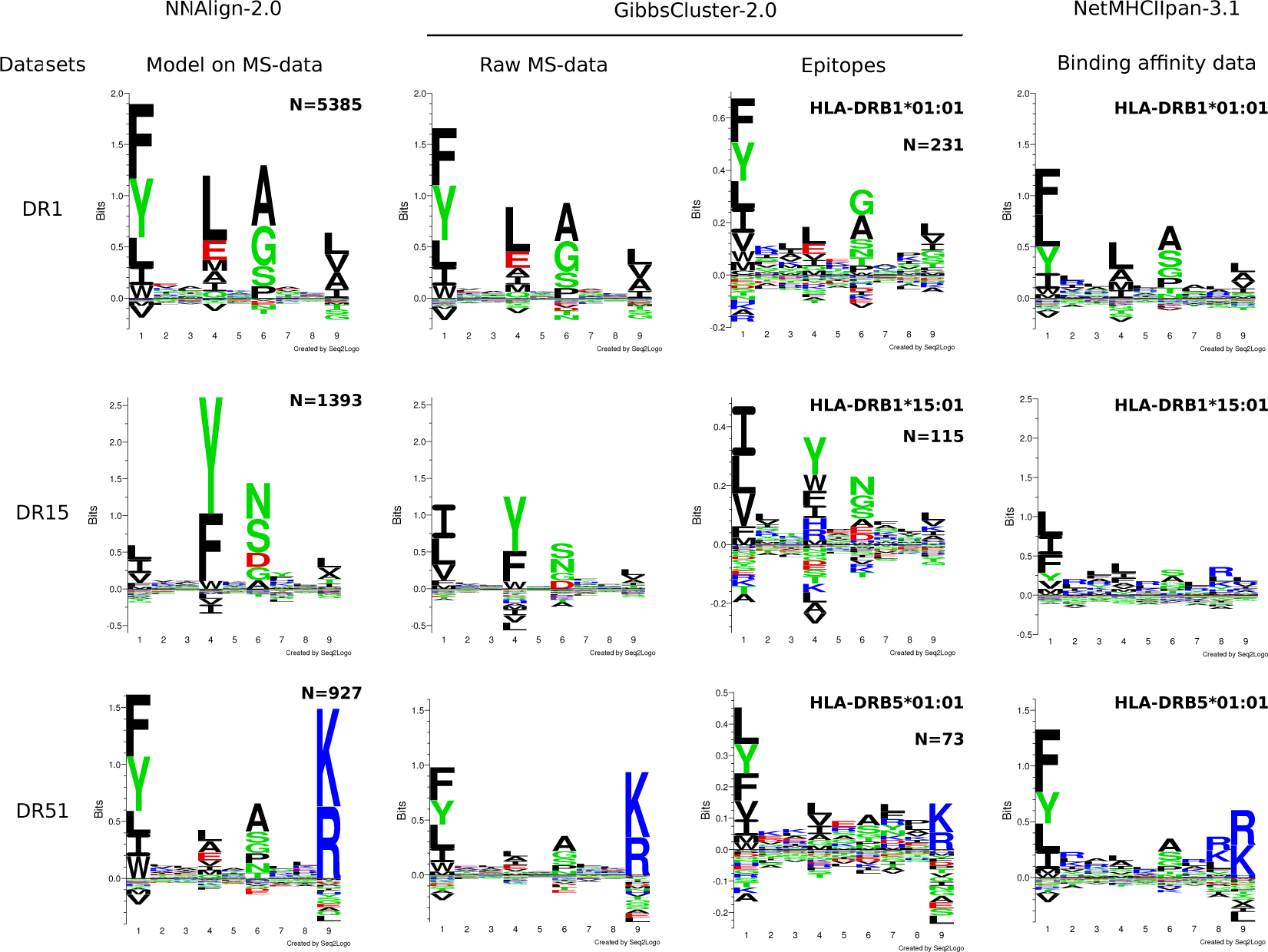
Comparison of motifs generated by different approaches for three HLA-DR alleles. NNAlign-2.0 motifs were obtained by training artificial neural networks on each MS data set, and evaluating 100,000 random peptides. The top scoring 1% peptides were used to build logos. Raw MS data were aligned, clustered and filtered in an unsupervised manner using GibbsCluster, with a trash cluster threshold = 2. The same procedure was applied to tetramer-positive data downloaded from the IEDB. Note that due to small data set size, epitope logos are shown in a different y-axis scale. Binding motifs for NetMHCIIpan-3.1 were determined by evaluating 100,000 random peptides, and visualizing the core motif of the top 1% scoring sequences.

## Final remarks

The binding specificities of MHC molecules have been traditionally characterized using *in-vitro* assays of binding affinity. The peptide-MHC binding data amassed through decades of painstakingly low-throughput experiments has had a tremendous contribution to the characterization of the binding preference for the most prevalent MHC molecules, and more generally to the understanding of the peptide repertoire available for T cell recognition. However, because of the extreme polymorphism of the MHC-encoding genes, with up to several thousand allelic variants per locus, the full characterization of their specificities remains infeasible. Tandem mass-spectrometry has emerged in the past decade has a powerful, high-throughput alternative for the identification of peptides eluted on the surface of antigen-presenting cells. The appeal of MS-based techniques does not only reside in the sheer amount of ligand data that can be detected in a single experiment, but also in the fact that it should capture additional signals of antigen processing besides the binding affinity measurable by *in-vitro* assays. Accurate tools for the identification of sequence motifs in eluted ligand datasets are essential to interpret the patterns underlying the immunopeptidome and to benefit from this data deluge.

In this report, we described some straightforward, efficient approaches to extract motifs from immunopeptidomes in a number of scenarios commonly encountered in the field. We outlined analyses for MHC class I and class II, both in cell lines expressing a single MHC allele and in unmodified cells with multiple MHC allelic variants. GibbsCluster [17] is our tool of choice because it can effectively remove residual contaminants after FDR filtering, deconvolute multiple motifs in a mixture of peptides of variable length and because it works both for MHC class I and class II ligands. In general, MHC class I molecules have strong, well-defined motifs, and even in samples containing several specificities it is often feasible to separate them into individual clusters. An unresolved problem remains the unambiguous association of each cluster to individual MHC molecules, especially for alleles with unknown binding motifs. So far only Bassani et al.

[25] have attempted to tackle this question, exploiting the co-occurrence of MHC class I alleles across different data sets of known haplotype to assign motifs to individual alleles. More work along these lines is needed to automatically annotate the MHC restriction of peptides in poly-allelic datasets. Current methods also assume that each peptide is restricted to one and only one MHC molecule. When cells express different alleles with similar binding motifs, or in the case of MHC class II ligands binding to multiple alleles in different alignment frames, it is likely that an individual peptide can act as ligand for multiple MHCs in a mixture. Future improvements to existing algorithm should aim to address this limitation and account for potential multiple restrictions of individual ligands.

A number of recent reports have described the first prediction methods trained directly on MHC class I ligand elution data from MS [3,4,26,45]. Their results indicate that methods trained on naturally presented peptides largely outperform prediction methods trained solely on *in-vitro* binding affinity data when it comes to identification of MHC ligands and epitopes. No reports have yet been published on models directly trained on MHC class II eluted ligands. Because the performance of MHC class II prediction methods still lags far behind their class I counterparts for epitope prediction, antigen processing factors are likely to play a major role in the generation of MHC class II ligands. Incorporating naturally processed ligand from MS experiments in the training pipelines of MHC class II prediction methods is an exciting and yet unexplored opportunity to close that gap. A simple but powerful approach to generate prediction models from ligand data is the NNAlign method [30]. We illustrate the construction of models from MS eluted ligands both for MHC class I and MHC class II, and show that they capture the preferences of the training data both in terms of binding motif and ligand length distribution. Taken together, these computational tools allow researchers to interpret motifs contained in immunopeptidomes and generate prediction models to scan protein databases for epitope candidates.

## Acknowledgements

This work was supported by Federal funds from the National Institute of Allergy and Infectious Diseases, National Institutes of Health, Department of Health and Human Services, under Contract No. HHSN272201200010C; and by the Agencia Nacional de Promoción Científica y Tecnológica, Argentina (PICT-2016-0089).

The authors have declared no conflict of interest.

## Supplementary Figures

**Supplementary Figure S1.**
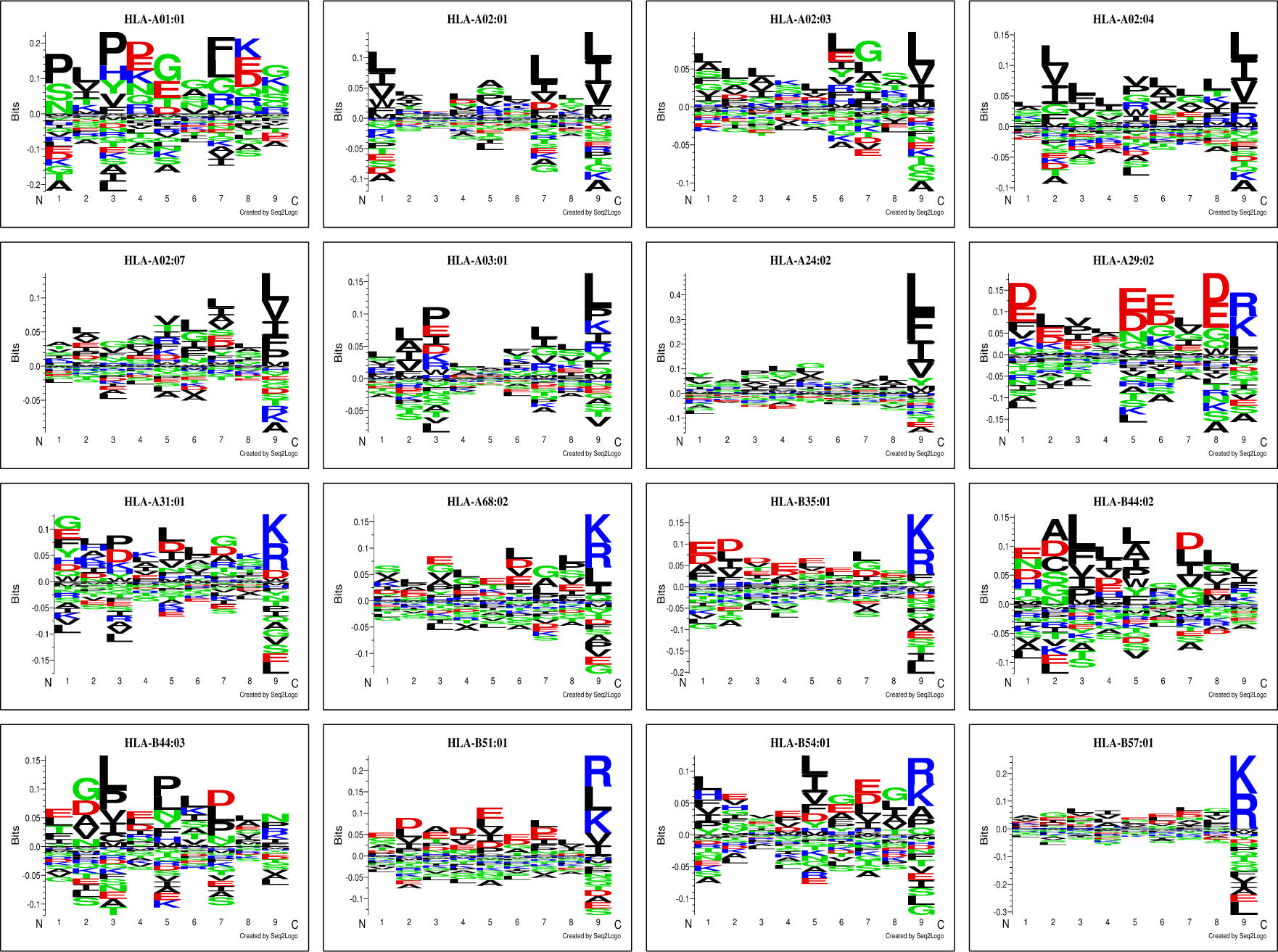
Sequence motifs of peptides collected by the trash cluster on the 16 alleles in the Abelin data set.

**Supplementary Figure S2.**
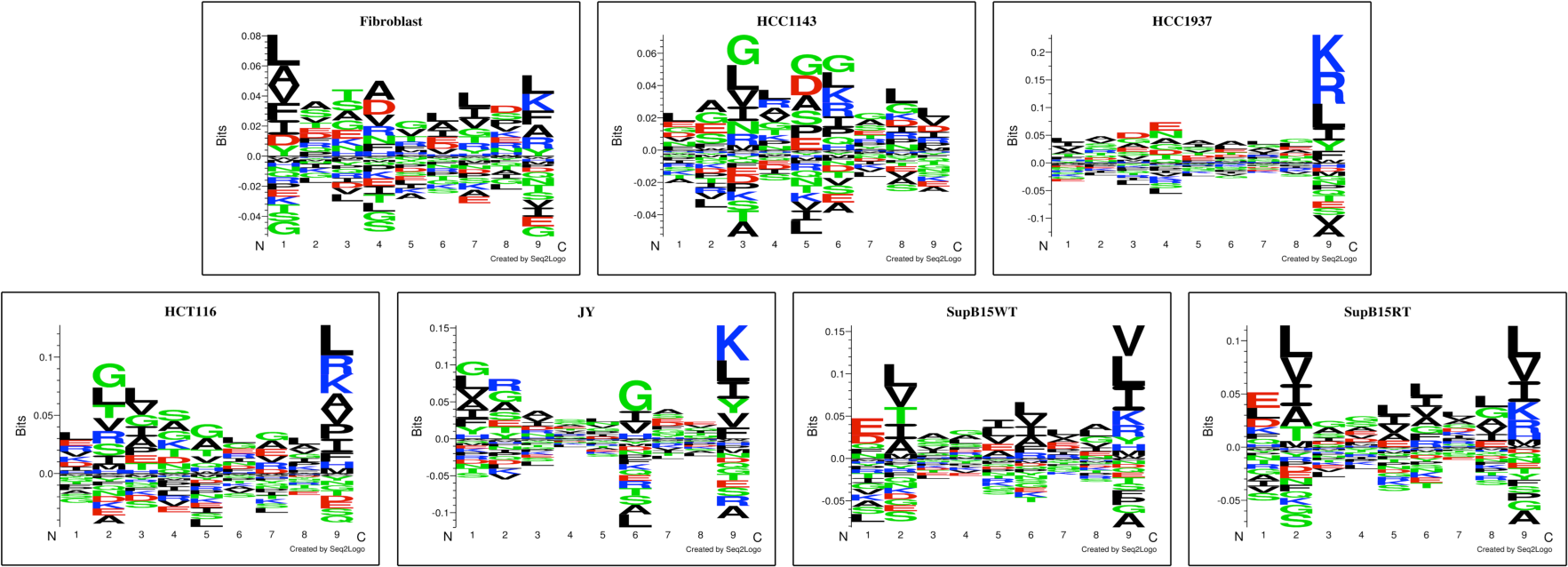
Sequence motifs of peptides collected by the trash cluster on the 7 alleles in the Bassani-Sternberg data set.

